# A phenomenological model of the time course of maximal voluntary isometric contraction force for optimization of complex loading schemes

**DOI:** 10.1101/256578

**Authors:** Johannes L. Herold, Christian Kirches, Johannes P. Schlöder

## Abstract

We construct a simple and predictive ordinary differential equation model to describe the time course of maximal voluntary isometric contraction (MVIC) force during voluntary isometric contractions and at rest. These time courses are of particular interest whenever force capacities are a limiting factor, e.g. during heavy manual work or resistance training (RT) sessions. Our model is able to describe MVIC force under complex loading schemes and is validated with a comprehensive set of published data from the elbow flexors. We use the calibrated model to analyze fatigue and recovery patterns observed in the literature. Due to the model’s structure, it can be efficiently employed to optimize complex loading schemes. We demonstrate this by computing a work-rest schedule that minimizes fatigue and an optimal isometric RT session as examples.

## 1 Introduction

Muscle fatigue is defined as an exercise-induced reduction in the ability to generate force or power [24]. Multiple task-specific mechanisms contribute to this complex phenomenon [19]. These mechanisms are categorized as peripheral (arising distal from the neuromuscular junction) or central (originating at spinal or supraspinal level). For a comprehensive overview of the physiological mechanisms, we refer to the work of Allen et al. [1] and Gandevia [24].

Several criteria can be used to measure muscle fatigue, e.g. the maximal voluntary isometric contraction (MVIC) force, power, the rate of force development, the one-repetition-maximum, or repetition strength [73, 57, 9]. To distinguish between peripheral and central contributions, one can superimpose external stimuli during MVICs either electrically to the nerve trunk or to the muscle belly [63], or magnetically to the motor cortex [72]. The resulting force increment then provides information about the origin of fatigue. During sub-maximal contractions and at rest, MVICs combined with external stimuli have to be interspersed to determine the corresponding time courses. For a review of these time courses, we refer to the work of Carroll et al. [13].

Predicting the time course of muscle fatigue and recovery would enable an optimal use of the limited human force capacities. With a suitable mathematical model, work shifts could be scheduled to avoid fatigue related accidents [29, 74] or resistance training (RT) sessions could be individualized to optimize adaptations [21]. Owing to the large number of possible combinations of contraction intensities and durations, these optimization problems are inherently high-dimensional. Thus, derivative-based methods have to be employed for an efficient solution, which imposes several mathematical requirements on model candidates.

This work deals with modeling the time course of MVIC force for several reasons. First, MVIC force is considered one of the gold standards for determining muscle fatigue [73], as its measurements are easy to standardize and to conduct. It is therefore used extensively by physiologists and sports scientists, who could benefit from a predictive model when designing experiments. Second, although most everyday movements are of a dynamic nature, isometric contractions contribute to stabilization during those movements [2]. For this reason, isometric strength capacities and especially their connection to possible injuries are of high interest, e.g. in ergonomics [36, 28] or sports [41]. Third, isometric strength is an important physical characteristic for a variety of athletes, e.g. wrestlers, gymnasts [68], or climbers [21]. Due to the specificity of adaptations, instead of dynamic exercises, isometric training is favorable for these athletes. Finally, isometric resistance training can be used as part of a rehabilitation program, e.g. when joint movements are restricted or not advisable [38].

### 1.1 Contributions

In this article, we provide a literature overview of mathematical and computational models describing the time course of MVIC force. We demonstrate that a new model is necessary as existing models suitable for derivative-based optimization do not perform satisfactorily. Based on physiological observations and existing work by Freund and Takala [22], we propose a phenomenological ordinary differential equation (ODE) model of the time course of MVIC force during voluntary isometric contractions and at rest. We use a comprehensive set of published data to validate the model for the elbow flexors and the model shows promising results. Furthermore, we illustrate the benefits of the model structure by computing a work-rest schedule that minimizes fatigue and an optimal isometric RT session as examples.

### 1.2 Structure of the article

The remainder of this article is structured as follows. In Section 2, we describe the data used for our work. In Section 3, we provide an overview of existing models and evaluate them with respect to our purposes. In Section 4, we justify the need for a new model. In Section 5, we present our model and validate it by estimating parameters from a subset of the available data and using those estimates to predict the remaining data. In Section 6, we use the calibrated model to examine fatigue and recovery patterns observed in the literature. Furthermore, we compute a work-rest schedule that minimizes fatigue and plan an optimal isometric RT session. In Section 7, we conclude with an outlook on possible extensions and applications.

## 2 Description of the data used

To the best of our knowledge, there is no study that reports time courses of MVIC force for the same muscle group of a single subject under different loading schemes. Thus, we use the mean values from several experiments examining muscle fatigue and recovery of the elbow flexors to evaluate the performance of the models under consideration.

The following experiments are used:

**E1** Taylor et al. [69] examine a 2 min MVIC and a recovery period lasting roughly 7 min for 8 subjects. Up to 30 MVIC force measurements are given per subject.

**E2** Søgaard et al. [66] examine a submaximal contraction at 15 % of baseline MVIC force lasting 43 min and 23 min of recovery for 9 subjects. 29 MVIC force measurements are given per subject.

**E3a** Taylor et al. [70] examine intermittent MVICs of 5 sec contraction and 5 sec rest and a 2 min recovery period for 9 subjects. Up to 18 MVIC force measurements are given per subject.

**E3b** Taylor et al. [70] examine intermittent MVICs of 15 sec contraction and 10 sec rest and a 4 min recovery period for 9 subjects. Up to 30 MVIC force measurements are given per subject.

**E3c** Taylor et al. [70] examine intermittent MVICs of 15 sec contraction and 5 sec rest and a 4 min recovery period for 8 subjects. Up to 30 MVIC force measurements are given per subject.

**E3d** Taylor et al. [70] examine intermittent MVICs of 30 sec contraction and 5 sec rest and a 3 min recovery period for 9 subjects. Up to 17 MVIC force measurements are given per subject.

**E4** Gandevia et al. [25] examine a 2 min MVIC and a subsequent 3 min recovery period for 8 subjects. 13 mean MVIC force values and standard errors of the means were extracted from the figures with the software Engauge Digitizer 10.0 [50].

**E5** Smith et al. [65] examine a 70 min submaximal contraction at 5 % of baseline MVIC force and the following 29 min of recovery for 8 subjects. 52 MVIC force measurements are given per subject. As the measurement times are only given for one subject, we use those for the whole sample.

All experiments employed a similar setup (i.e. elbow flexed to 90 degrees, forearm vertical and supinated [69, 66, 70, 25, 65]), which justifies a comparison of these values. To the best of our knowledge, this is the most comprehensive set of data used to validate a model of the time course of MVIC force.

For each experiment *k* ∈ {E1, E2, E3a, E3b, E3c, E3d, E4, E5}, we are using the mean values

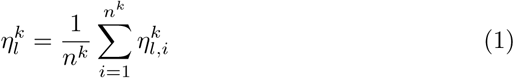

of the *n*^*k*^ individual measurements 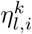 to validate our model. The measurement errors 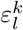 are assumed to be additive, independent, and identically normally distributed with mean zero and standard deviation 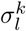. We take 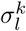 to be the corrected sample standard deviation (SD) of the corresponding mean value 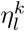 divided by the square root of the sample size *n*^*k*^, i.e.

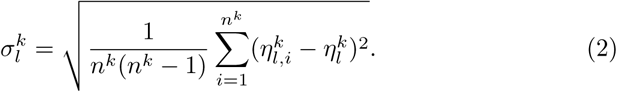

Thus, 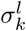 yields an approximation of the standard error of the mean. For experiment E4, the means and the corresponding standard errors were extracted directly from the figures. The test contractions interspersed during submaximal contractions and recovery are described to last 1-2 [70] or 2 - 3 sec [66]. Therefore, we model those contractions to last 2 sec.

**Remark**. Since we are working with isometric contractions, force and torque normalized to baseline are equivalent. Thus, we do not differentiate between the two terms in the following.

## 3 Literature overview

Several mathematical and computational models have been proposed to predict the time course of MVIC force during voluntary isometric contractions or the time course of maximal evocable isometric force under external stimulation. As fatigue is highly task-dependent and substantial differences exist between evoked and voluntary contractions [48], we do not include models created for external stimulation in the following overview. Furthermore, we assume that all models can be applied at muscle group level and account for peripheral and central factors contributing to MVIC force generation.

### 3.1 Evaluation criteria

We put a special focus on the following criteria necessary for our intentions.

**C1** Can the model predict fatigue of MVIC force under complex patterns of voluntary isometric contractions, i.e. maximal, submaximal, sustained, and intermittent contractions of varying intensity?

**C2** Does the model include recovery of MVIC force?

**C3** Is the model suitable for real-life applications, i.e. is the number of parameters low and are all parameters identifiable through MVIC force measurements?

**C4** Is the model suitable for high-dimensional optimization, i.e. is the model suitable for derivative-based solution methods?

Many physiology-based models implement a feedback loop with some kind of proportional-integral-derivative controller to imitate force adjustments by the nervous system and match simulated to target force. Unfortunately, these closed-loop controllers violate criterion C4 as they are usually not suitable for derivative-based optimization methods without considerable effort spent on reformulations.

### 3.2 Existing models

First attempts to quantify the development of muscle fatigue were made around 1960 by Monod and Scherrer [51, 52] and Rohmert [58], who mathematically described the nonlinear relationship between contraction intensity and endurance time with algebraic equations. Subsequently, several authors proposed similar equations or developed joint-specific versions [18, 40]. However, these models neither allow to evaluate the time course of MVIC force for more complex contraction scenarios nor do they include recovery.

Fuglevand et al. [23] constructed a model of recruitment and rate coding of a pool of motor units. As the model does not include fatigue effects, it was later on extended by Dideriksen et al. [17]. The fatigue-induced changes of the model require a controller to maintain target force. Potvin and Fuglevand [53] proposed another modification to the original model [23], which has yet to be extended to be able to describe force recovery.

Hawkins and Hull [30] incorporated fatigue effects into a previously developed fiber based model [31]. The model does not describe recovery after work and employs an if-else structure to account for the recruitment order of different fiber types.

Based on the work of Rich [55] and Deeb et al. [16], Wood et al. [74] proposed a model and subsequently used it to minimize fatigue during work schedules consisting of intermittent contractions with constant intensities and duty cycles. The least fatiguing setting was found by an exhaustive search through the parameter space. Although their approach pursued a similar goal as this work, the fixed duty cycles impose an undesired limitation on the loading schemes.

Freund and Takala [22] constructed a biomechanical model of the forearm and introduced a simple ODE to account for effects of fatigue and recovery. Ma et al. [46, 47] utilized similar dynamics but separated fatigue and recovery, which resulted in a branchwise definition of the ODE’s right-hand-side. Riener et al. [56] modeled the effects of fatigue and recovery for externally stimulated contractions. Although they chose muscle activation as model input, the proposed dynamics are closely related to the ones of Freund and Takala [22] and Ma et al. [47], which is why we mention it here. Fayazi et al. [20] based their model on that of Ma et al. [47], but modeled fatigue and recovery to occur simultaneously as originally proposed by Freund and Takala [22] for voluntary contractions and by Riener et al. [56] for stimulated contractions. Fayazi et al. [20] used their model subsequently to calculate optimal pacing strategies for a cyclist. The type of model proposed by these authors fulfills our requirements and is thus examined further in the next section.

Liu et al. [44] introduced a three-compartment model distinguishing between active, fatigued, and resting motor-units. Fatigue and recovery effects are represented by flows between the compartments and brain effort was chosen as model input. Since brain effort is only known for maximal efforts, the model was later extended by Xia and Law [75] by including a controller to account for the recruitment hierarchy of three different fiber types when matching the target force. A different modification of the original model [44] was implemented by Sih et al. [64]. The authors expanded the model to four compartments and reformulated their equations to circumvent brain effort as input. In general, this allows the model to simulate arbitrary force profiles. Nevertheless, it is still defined branchwise.

Several authors have extended or modified the model of Xia and Law [75]. Gede and Hubbard [27] added a force-velocity dependence and generalized the model to task level. Furthermore, they removed the fast components of the model and changed the input to be the amount of active muscle which makes the model suitable for optimization [26]. However, because of the separate inputs for each fiber type, the model could only be validated for maximal efforts. Other modifications to the original model [75], e.g. using time-variant [67] or contraction-specific [45] parameters, have been developed as well.

James and Green [34] used a similar approach as Sih et al. [64] but assumed that contractile properties vary as continuous functions of time and motor unit type. The power output of a single motor unit is defined non-smooth and the model does not account for force recovery. Callahan et al. [10, 11] built a comprehensive model of torque generation. The model can be used to describe voluntary and externally stimulated contractions and uses a controller to match generated to target torque. Contessa and Luca [15] developed a model focusing on motor units containing a feedback loop implemented to regulate the excitation and match target force.

## 4 Evaluation of suitable models

As discussed in the previous section, the models proposed by Freund and Takala [22] and Fayazi et al. [20] are suitable for our intentions. Thus, we present these models in detail and evaluate their performance by fitting them to the available data.

### 4.1 Freund and Takala model

Freund and Takala [22] modeled fatigue and recovery of MVIC force as

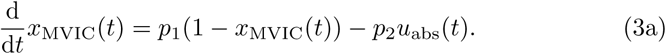

Here,

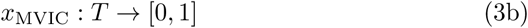

represents the current MVIC force capacity on the time interval *T* ⊂ [0, ∞) and

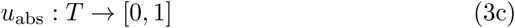

represents the absolute external isometric load and is the model input. The fatigue rate *p*_2_*u*_abs_(*t*) depends linearly on *u*_abs_. The recovery rate *p*_1_(1–*x*_MVIC_(*t*)) depends linearly on the difference between the current force capacity *x*_MVIC_ and the maximal force capacity. Since we scale all states and inputs to [0,1], the maximal force capacity equals 1. The dimensionless parameters *p*_1_ ∈ [0, ∞) and *p*_2_ ∈ [0, ∞) determine the maximal speed of exponential recovery and fatigability of the muscle group. The initial condition

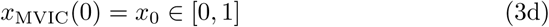

describes the force capacity of the muscle group at *t* = 0. For an unfatigued muscle, we choose *x*_0_ = 1.

### 4.2 Fayazi et al. model

Based on the work of Ma et al. [47], Fayazi et al. [20] proposed dynamics similar to those of Freund and Takala [22]. Keeping the notation introduced earlier, the dynamics of the model read as

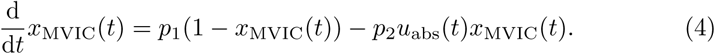

Thus, the difference to the model of Freund and Takala [22] is an additional dependency of the fatigue rate on the current MVIC force. In the model of Ma et al. [47], where fatigue and recovery are modeled separately, this factor ensures a sufficient decrease of fatigability during an MVIC.

### 4.3 Simulating MVICs

To simulate maximal voluntary contractions, it is favorable to substitute

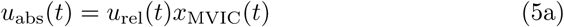

and use

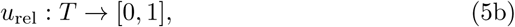

the load relative to the current force capacity, as input. This substitution reflects the experimental settings of an MVIC, during which subjects are asked to contract maximally (*u*_rel_ = 1) instead of maintaining a certain target force *u*_abs_. It furthermore allows us to simulate MVICs without an unnecessarily complex mathematical description of the non-constant input force *u*_abs_ and is necessary to model MVICs for which no measurement values are given (e.g. in experiment E3a).

However, in contrast to using *u*_abs_ as input, this substitution only gives a prediction of the absolute external load at a certain level of effort *u*_rel_ by the model and not the actual absolute external load. Depending on whether the contraction is submaximal or maximal, we either use *u*_abs_ or *u*_rel_ to describe the experimental settings in the following.

### 4.4 Necessity of a new model

We now fit these models simultaneously to the data of E1 and E2 presented in Section 2. The multiple experiment parameter estimation problem for one model is formulated as

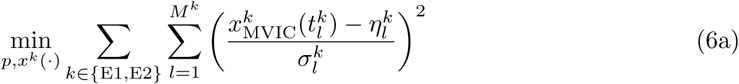

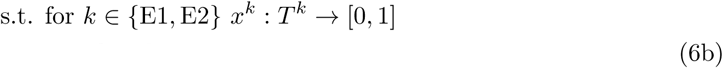

satisfies the model equations (3) or respectively (4)

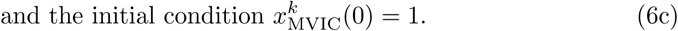

Here, *M*^*k*^ ∈ ℕ is the number of measurements in experiment *k* ∈ {E1, E2} and the *l*-th measurement 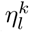 of experiment *k* is characterized by its time 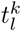 and the standard deviation 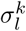 of its measurement error. As explained previously, the loading scheme of experiment *k* is described by the input function *u*_abs_ or via substitution by *u_rel_*. The state 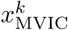 then denotes the model response for this setting.

The optimization problem constrained by an ODE (6) is solved by employing a first-discretize-then-optimize strategy. We use a direct multiple shooting approach to reduce the problem to a finite-dimensional form [7] and employ DAESOL [4] for integration of the ODE and sensitivity generation via internal numerical differentiation [6]. Afterwards, the resulting structured nonlinear least squares problem is solved with PAREMERA [37], an implementation of the reduced generalized Gauss-Newton method [62]. Both packages are embedded in the optimal experimental design software VPLAN [39].

Figure 1 illustrates the results of the fit for the model of Freund and Takala [22] and Figure 2 for the model of Fayazi et al. [20]. Table 1 lists the parameter estimates and their estimated relative standard deviations for both models. Table 2 lists the mean absolute errors and weighted residual sum of squares of the fits for both models.

**Figure 1:**
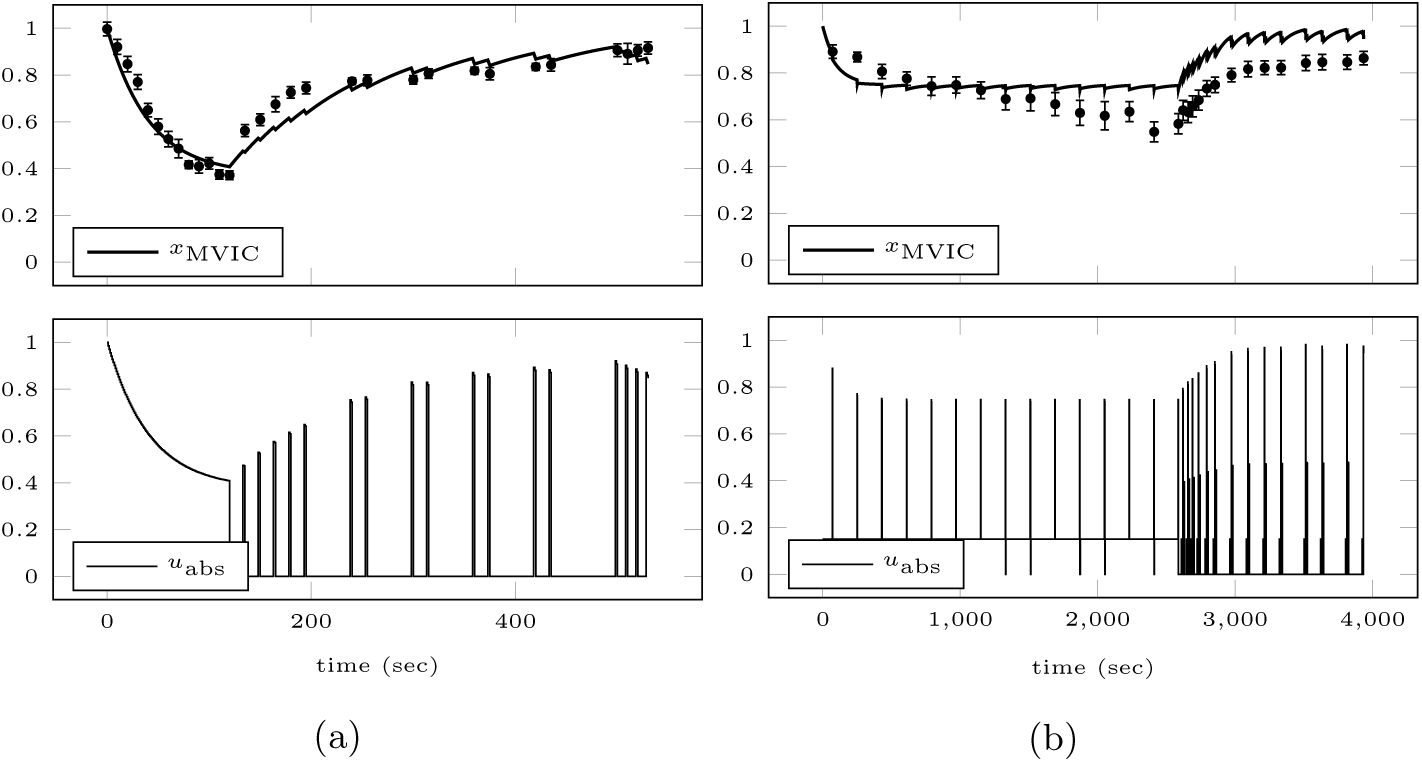
Model response obtained by fitting the model of Freund and Takala [22] simultaneously to the data of E1 (a) and E2 (b). The top row shows the mean values of the experiments plotted against the model response for the experimental setting. The error bars represent the standard errors of the means. Additionally, the absolute force input as predicted by the model (explained in Section 4.3) is illustrated in the bottom row.

**Figure 2:**
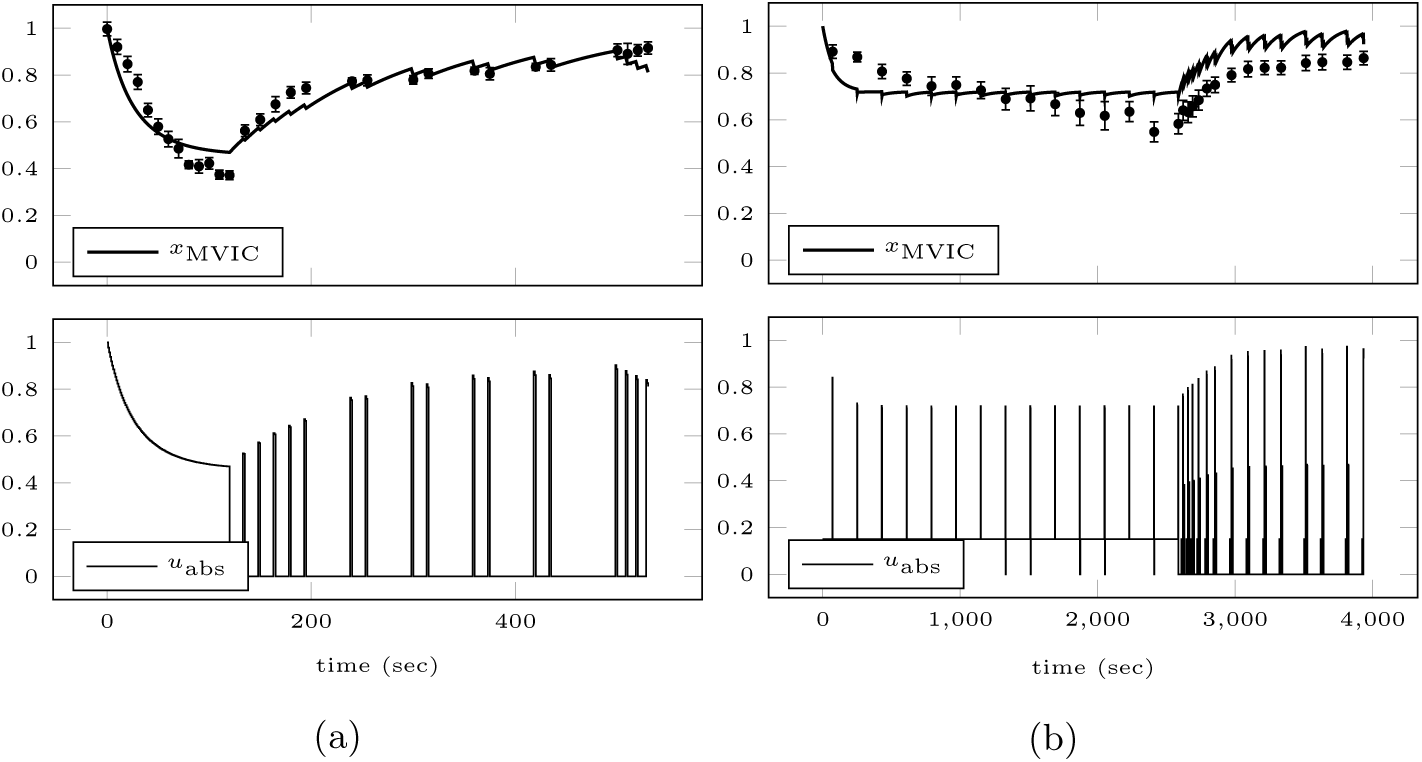
Model response obtained by fitting the model of Fayazi et al. [20] simultaneously to the data of E1 (a) and E2 (b). The top row shows the mean values of the experiments plotted against the model response for the experimental setting. The error bars represent the standard errors of the means. Additionally, the absolute force input as predicted by the model (explained in Section 4.3) is illustrated in the bottom row.

**Table 1:** Parameter estimates and their estimated relative standard deviations obtained by simultaneously fitting the models of Freund and Takala [22] and Fayazi et al. [20] to the data of E1 and E2.

| | $p_1$ | ± SD | $p_2$ | ± SD |
| --- | --- | --- | --- | --- |
| Freund and Takala [22] | $9.20 \cdot 10^{-3}$ | $\pm 3.75 \%$ | $1.53 \cdot 10^{-2}$ | $\pm 3.56 \%$ |
| Fayazi et al. [20] | $8.66 \cdot 10^{-3}$ | $\pm 4.90 \%$ | $2.24 \cdot 10^{-2}$ | $\pm 4.98 \%$ |

**Table 2:** Mean absolute errors (MAE) and weighted residual sum of squares (WRSS) obtained by simultaneously fitting the models of Freund and Takala [22] and Fayazi et al. [20] to the data of E1 and E2.

|  | E1 |  | E2 |  |
| --- | --- | --- | --- | --- |
|  | MAE | WRSS | MAE | WRSS |
| Freund and Takala [22] | 0.04 | 151.07 | 0.10 | 273.72 |
| Fayazi et al. [20] | 0.05 | 210.97 | 0.09 | 234.80 |

When evaluating the fits, the necessity of a new model becomes clear. The model of Freund and Takala [22] captures the fatigue development during the MVIC of E1 correctly, whereas the model of Fayazi et al. [20] underestimates it slightly. This is caused by combining the fatigue and recovery branches of Ma et al. [47] linearly. Both models overestimate fatigue in the beginning of the submaximal contraction E2 and reach a steady-state too early. Furthermore, due to their mono-exponential recovery terms, they cannot capture the initially faster recovery after cessation of the contractions.

## 5 Proposed model

In this section, we propose a new ODE model of the time course of MVIC force, motivate our choice, and validate the model.

### 5.1 Structure of the model

As the model of Freund and Takala [22] performed slightly better in the previous fits, we use their model as a foundation. The proposed new model is given as

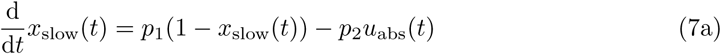

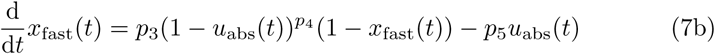

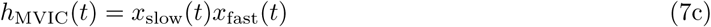

where

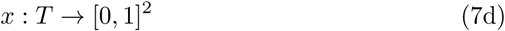

is defined on the time horizon *T* ⊂ [0, ∞) and consists of two state variables *x*_fast_ and *x*_slow_. Again, we scale all states and inputs to [0,1]. The model furthermore contains five dimensionless parameters *p*_*j*_ ∈ [0, ∞) for *j* ∈ {1, …, 5} and one input function

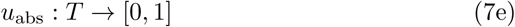

representing the absolute external isometric load. The current MVIC force capacity is denoted by

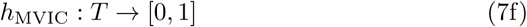

and the initial conditions for the states are given by

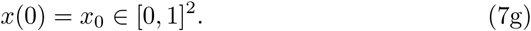

For an unfatigued muscle, one chooses *x*_0_ = (1, 1)^*T*^.

Table 3 summarizes the model components. To increase readability throughout the rest of this paper, we omit arguments of the states and input functions whenever the dependencies are clear.

**Table 3:** Overview of the model components.

|  | Type | Interpretation |
| --- | --- | --- |
| $x_{\text{slow}}$ | State variable | Slow component |
| $x_{\text{fast}}$ | State variable | Fast component |
| $h_{\text{MVIC}}$ | Function | MVIC force |
| $p_1$ | Parameter | Maximal recovery rate of $x_{\text{slow}}$ |
| $p_2$ | Parameter | Maximal fatigue rate of $x_{\text{slow}}$ |
| $p_3$ | Parameter | Maximal recovery rate of $x_{\text{fast}}$ |
| $p_4$ | Parameter | Influence of $u_{\text{abs}}$ on recovery rate of $x_{\text{fast}}$ |
| $p_5$ | Parameter | Maximal fatigue rate of $x_{\text{fast}}$ |
| $u_{\text{abs}}$ | Input function | Absolute external load |

### 5.2 Motivation for choosing this structure

We duplicated the dynamics of Freund and Takala [22] and modeled MVIC force as the product of two state variables. This was motivated by the observations of fast and slow dynamics of fatigue and recovery [13] and furthermore supported by other authors who proposed using double-exponential functions to describe the time courses of fatigue [16] and recovery [14]. The product was chosen to represent the chain of mechanisms leading to force generation. Yet, since this chain includes complex feedforward and feedback mechanisms, the strict separation is obviously an oversimplification.

The processes summarized in *x*_slow_ play a bigger role in contractions of long duration or repeated intermittent ones, as this state fatigues much slower and takes longer to recover. Correlations to the Ca^2+^ sensitivity of the crossbrigdes, to the Ca^2+^ release of the sarcoplasmic reticulum [1], and to glycogen depletion [61] are likely. Furthermore, there is also a long lasting effect of reduced voluntary activation after prolonged contractions, the reason for which is currently unknown [13].

*x*_fast_ pools a variety of mechanisms responsible for a quick onset of MVIC force reduction and a fast recovery. These mechanisms contribute to a larger degree during maximal and near maximal contractions and seem to be closely related to muscle perfusion [13]. Possible factors include metabolite accumulation (e.g. inorganic phosphate) [1], firing of group III and IV afferents [71], or energy deficiency [61]. Since studies have established a link between blood occlusion in the muscle (either through high-intensity contractions or artificially) and impeded recovery [5, 35], we added a dependency of the recovery rate on (1 – *u*_abs_)^*p*_4_^. The exponent *p*_4_ determines the contraction intensity at which total occlusion occurs [3, 60]. A similar dependency on activation was proposed by Riener et al. [56] in the context of external stimulation. However, they assumed a linear relationship.

### 5.3 Fitting the model to the data

To validate our model, we fit it to the data of E1, E2, E3a, and E3c. This subset was chosen as it contains a sustained maximal, a sustained submaximal, and two intermittent maximal contractions and enables an identification of the parameters. For details on the multiple experiment parameter estimation problem, we refer to Section 4.4. Figure 3 depicts the results of the fit. Table 4 lists the parameter estimates and their estimated relative standard deviations. Table 5 lists the mean absolute errors and weighted residual sum of squares of the fits.

**Figure 3:**
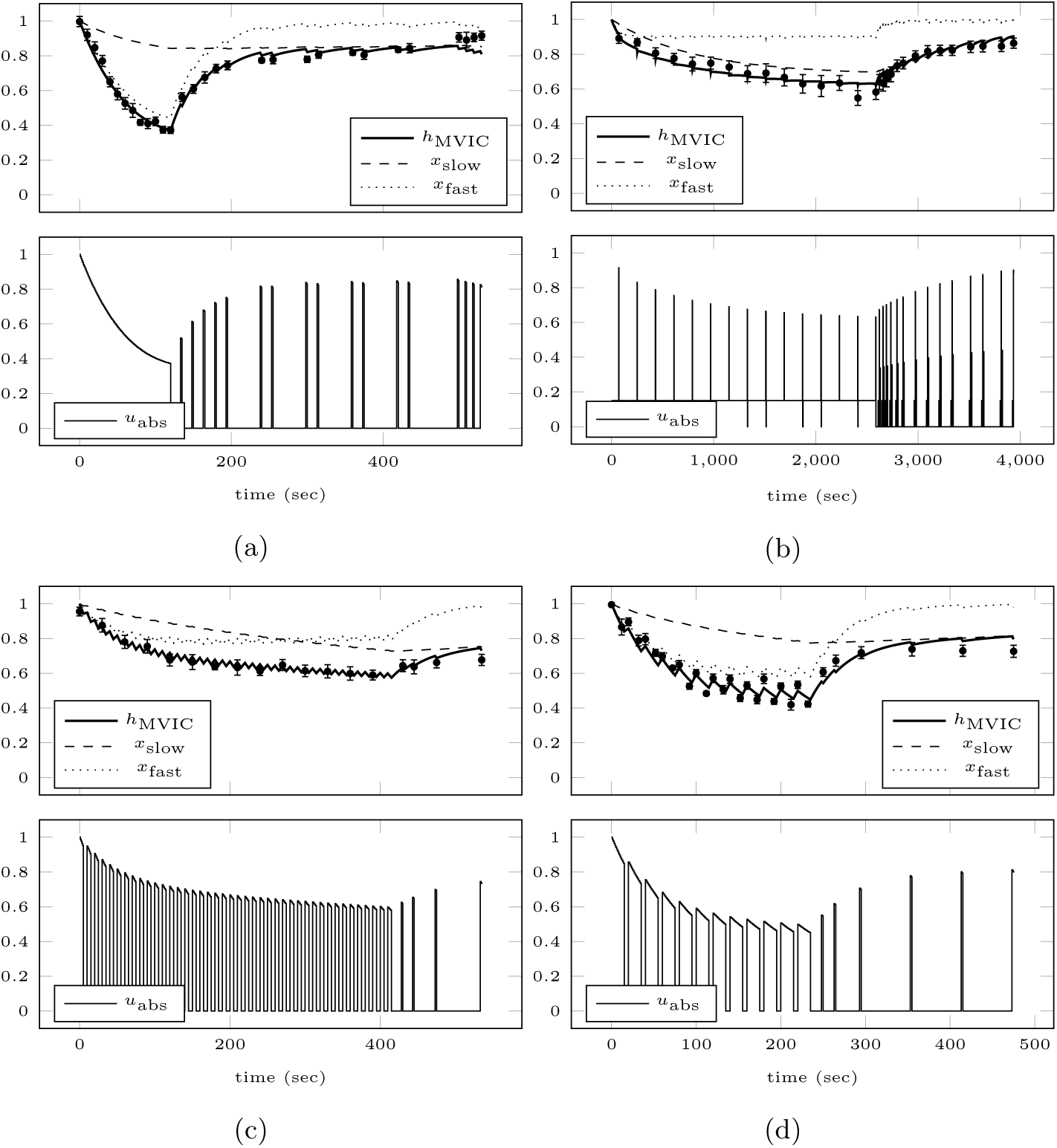
Model response obtained by fitting the proposed model simultaneously to the data of E1 (a), E2 (b), E3a (c), and E3c (d). The top row shows the mean values of the experiments plotted against the model response for the experimental setting. The error bars represent the standard errors of the means. Additionally, the absolute force input as predicted by the model (explained in Section 4.3) is illustrated in the bottom row.

**Table 4:** Parameter estimates and their estimated relative standard deviations obtained by simultaneously fitting the proposed model to the data of E1, E2, E3a, and E3c.

| | Interpretation | Estimate | ± SD |
| --- | --- | --- | --- |
| $p_1$ | Maximal recovery rate of $x_{\text{slow}}$ | $1.20 \cdot 10^{-3}$ | $\pm 7.23$ % |
| $p_2$ | Maximal fatigue rate of $x_{\text{slow}}$ | $2.46 \cdot 10^{-3}$ | $\pm 6.04$ % |
| $p_3$ | Maximal recovery rate of $x_{\text{fast}}$ | $2.85 \cdot 10^{-2}$ | $\pm 6.44$ % |
| $p_4$ | Influence of $u_{\text{abs}}$ on recovery rate of $x_{\text{fast}}$ | 3.99 | $\pm 11.84$ % |
| $p_5$ | Maximal fatigue rate of $x_{\text{fast}}$ | $9.44 \cdot 10^{-3}$ | $\pm 2.99$ % |

**Table 5:** Mean absolute errors (MAE) and weighted residual sum of squares (WRSS) obtained by simultaneously fitting the proposed model to the data of E1, E2, E3a, and E3c.

|  | E1 | E2 | E3a | E3c |
| --- | --- | --- | --- | --- |
| MAE | 0.03 | 0.03 | 0.02 | 0.03 |
| WRSS | 66.68 | 25.82 | 17.17 | 80.83 |

Although more data was used, the fit is improved substantially compared to Figures 1 and 2. This is to be expected, as the proposed model employs five instead of two parameters. Yet, since all parameters could be estimated reasonably well, the model is not overparameterized. Our model is able to capture submaximal contractions, the initial fast recovery after cessation of a contraction, and the delayed recovery after prolonged contractions.

Due to the phenomenological nature of the model, most of the estimated parameter values do not allow a direct physiological interpretation. Nevertheless, we expect physiological characteristics like fiber type composition, capillariza-tion, buffering capacity, muscle mass and strength, energy stores, and others to be reflected in the dimensionless parameter estimates. An inter-individual comparison of the estimates from the same muscle group or an intra-individual comparison of those from different muscle groups might provide interesting physiological insights. However, to evaluate this hypothesis, more data is needed.

Since the term (1 – *u*_abs_)^*p*_4_^ was specifically introduced to mimic blood occlusion during intense contractions, we compare the estimate of *p*_4_ to the values given in literature. Sadamoto et al. [60] determined that at a contraction intensity of 53 % of baseline MVIC force, muscle blood flow in the biceps was arrested. For the external load *u*_abs_ = 0.53, the estimated exponent *p*_4_ = 3.99 would imply roughly 5 % of the induced blood flow reaching the elbow flexors and providing some kind of recovery. As we are not only considering the biceps but all elbow flexors, and as Barnes [3] found a strength-dependency of the intensity necessary for total occlusion, this value seems to be consistent with the literature.

### 5.4 Testing the predictive ability of the model

The data of the remaining four experiments E3b, E3d, E4, and E5 is now used to test the predictive ability of the calibrated model. Figure 4 illustrates the prediction of the calibrated model for these experiments. Table 6 lists the mean absolute errors and weighted residual sum of squares of the predictions.

**Figure 4:**
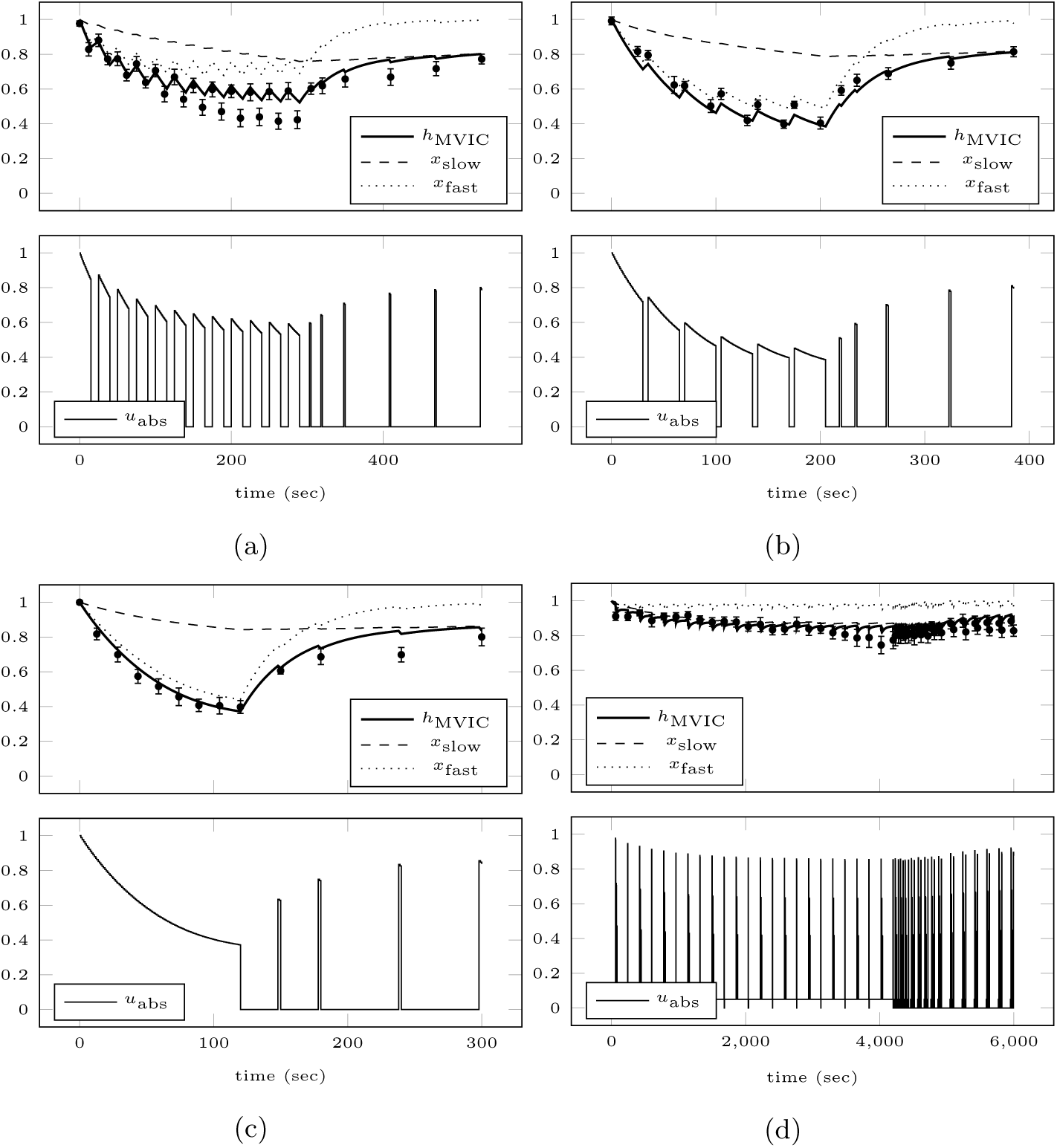
Model response obtained by simulating the calibrated model for the experimental settings of E3b (a), E3d (b), E4 (c), and E5 (d). The top row shows the mean values of the experiments plotted against the model response for the experimental setting. The error bars represent the standard errors of the means. Additionally, the absolute force input as predicted by the model (explained in Section 4.3) is illustrated in the bottom row.

**Table 6:** Mean absolute errors (MAE) and weighted residual sum of squares (WRSS) obtained by using the calibrated model to predict the data of E3b, E3d, E4, and E5.

|  | E3b | E3d | E4 | E5 |
| --- | --- | --- | --- | --- |
| MAE | 0.04 | 0.03 | 0.03 | 0.03 |
| WRSS | 47.53 | 37.86 | 15.28 | 41.00 |

Using the estimated parameters, the model is able to predict the data of E3d, E4, and E5 satisfactorily. Nevertheless, it shows unexpected deficiencies when simulating experiment E3b. Namely, it cannot capture the increasing fati-gability of the subjects during the experiment. The exact reasons are unknown. However, this increasing fatigability cannot be observed during the other experiments of Taylor et al. [70]. Furthermore, this increase actually contradicts the size principle [32] according to which primarily fatigue-resistant fibers should be used as the protocol progresses. Thus, fatigability should decrease during each contraction and a steady-state should be reached, as can be seen in the other experiments. Since this experiment additionally shows the largest inter-subject variability compared to other experiments from the study by Taylor et al. [70], we suppose that motivational issues of some subjects are the cause of this phenomenon. We argue similarly concerning the force drops during the submaximal contractions E2 and E5 shortly before the end of the contraction, which are not captured by the model.

## 6 Numerical case studies

In this section, we illustrate how the calibrated model can be used for analysis and optimization of real-world scenarios. Throughout this section, we use the parameter values of the elbow flexors obtained in Section 5.3.

### 6.1 Model-based analysis of fatigue and recovery patterns

We now examine two observations from the literature regarding the fatigue and recovery patterns of MVIC force. Since the model has been validated with contraction intensities of 15 % of baseline MVIC force and MVICs, we use those for our simulations. Furthermore, the plots in Section 5 illustrate that short test MVICs interspersed during submaximal contractions and recovery do not substantially influence the time course of MVIC force and can therefore be neglected in our simulations.

First, Rozand et al. [59] observe that after sustained isometric contractions of the knee extensors with different intensities and similar force-time-integral (FTI) the induced level of fatigue did not differ. The FTI on the time interval [0, *T*] is defined as

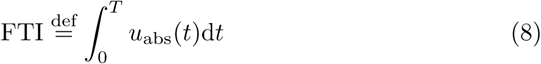

and often used as an analogue for work during isometric contractions, where no actual physical work is performed. Figure 5 shows the calibrated model simulated for a 60 sec MVIC and a submaximal contraction at 15 % of baseline MVIC force lasting 292.40 sec with the same FTI of 43.86. Both contractions end before a steady-state is reached. As can be seen in the plots, the induced fatigue is higher for the MVIC than for the submaximal contraction. After the maximal contraction, *h*_MVIC_ is reduced to 53.21 %. In contrast to this, *h*_MVIC_ is reduced to 82.37 % after the submaximal contraction. We therefore cannot reproduce the results of Rozand et al. [59] for our settings.

**Figure 5:**
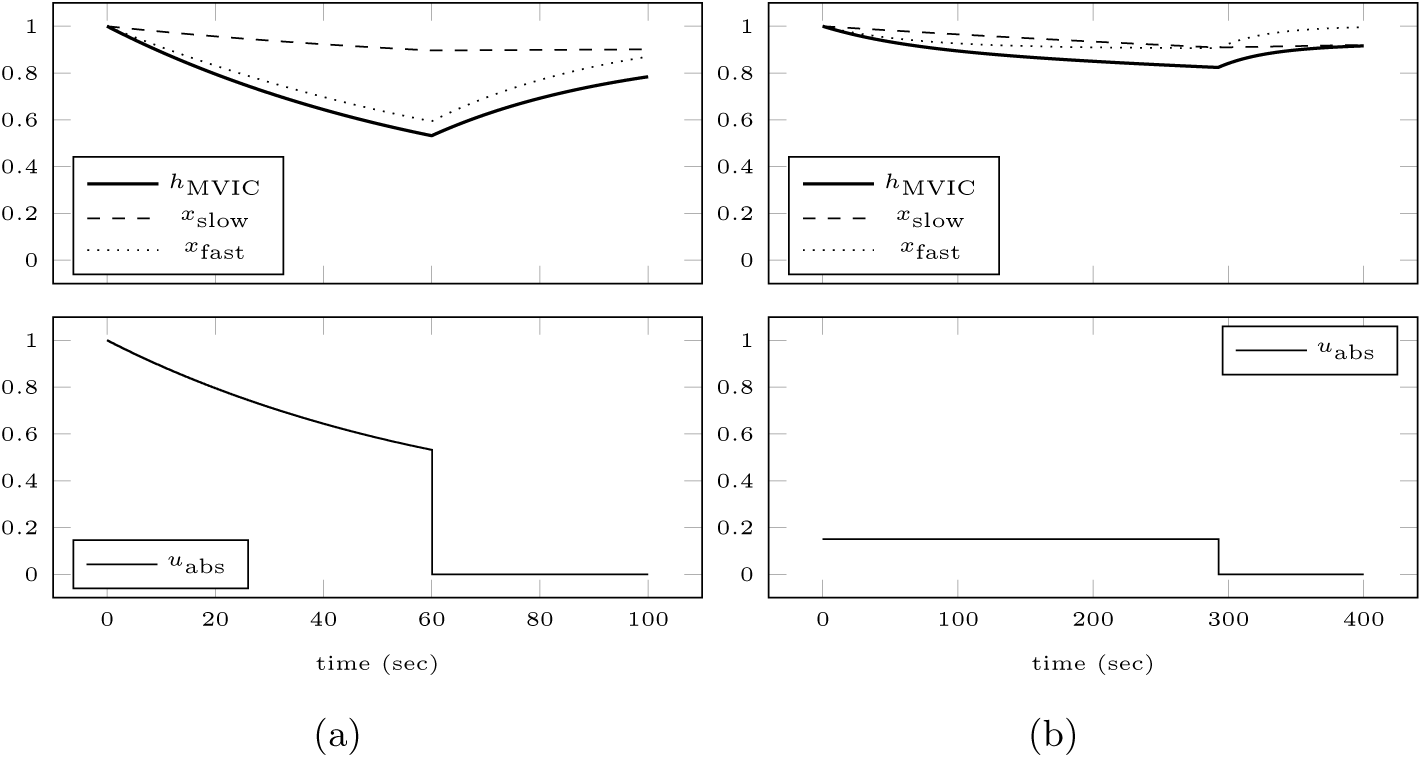
Model response obtained by simulating the calibrated model for an MVIC (a) and for a submaximal contraction (b) with the same force-time-integral. The top row shows the model response and the absolute force input is illustrated in the bottom row. The induced fatigue is higher for the MVIC than for the submaximal contraction.

Second, Rashedi and Nussbaum [54] observe that recovery is fatigue-and not task-dependent for intermittent isometric contractions of the index finger. Iguchi et al. [33] notice similar dependencies after sustained isometric contractions of the quadriceps. Figure 6 shows the calibrated model simulated for a 40 sec MVIC and a submaximal contraction at 15 % of baseline MVIC force lasting 2310 sec both reducing MVIC force to 64.41 % of baseline. Both contractions end before a steady-state is reached and are followed by a 10 min recovery period. As can be seen in the plots, recovery from the MVIC is almost complete after 10 min, whereas recovery from the submaximal contraction takes much longer. At the end of the MVIC scenario, *h*_MVIC_ recovered to 96.25 %. In contrast to this, *h*_MVIC_ recovered to 85.97 % at the end of the submaximal scenario. Thus, we cannot reproduce the results of Rashedi and Nussbaum [54] for our settings.

**Figure 6:**
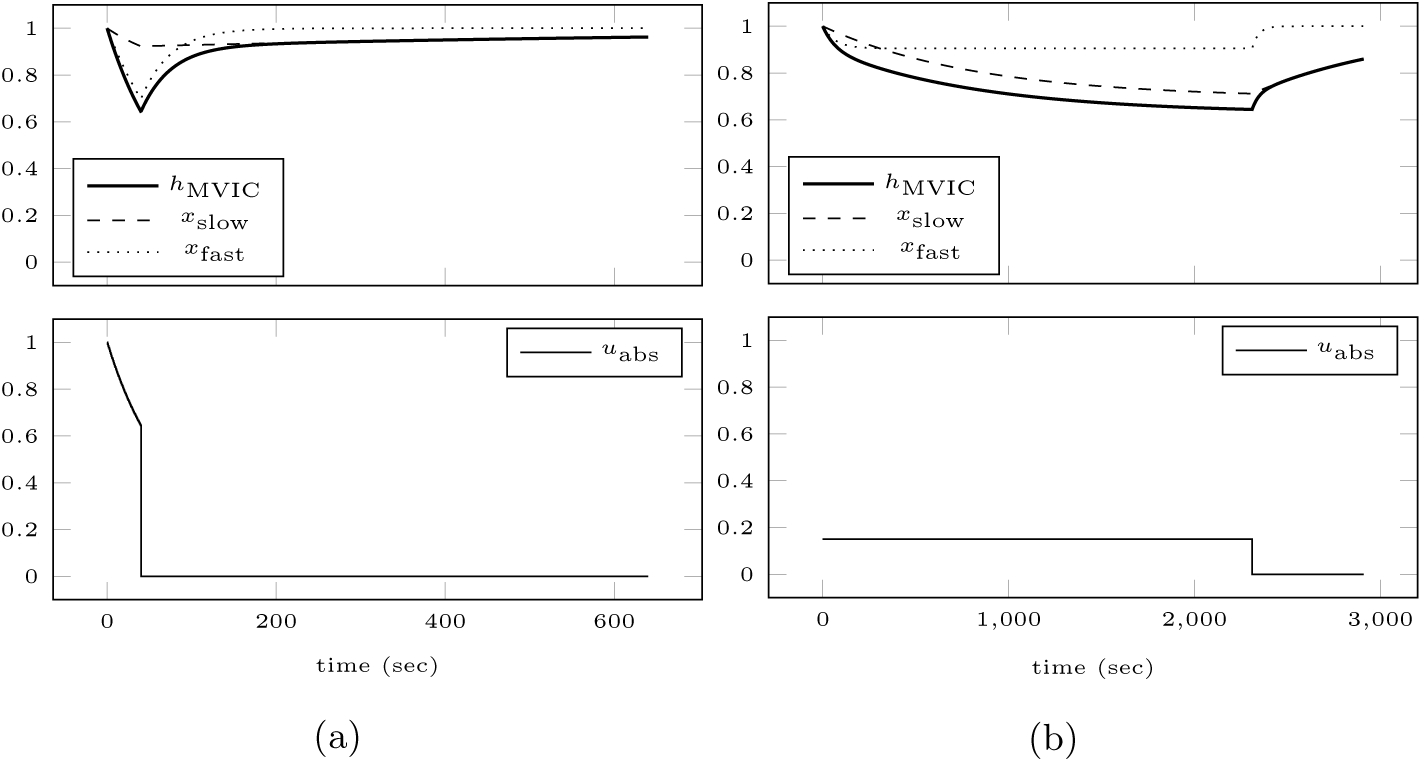
Model response obtained by simulating the calibrated model for an MVIC (a) and for a submaximal contraction (b) inducing the same level of fatigue followed by 10 min of recovery. The top row shows the model response and the absolute force input is illustrated in the bottom row. Recovery from the MVIC is almost complete after 10 min, whereas recovery from the submaximal contraction takes much longer.

### 6.2 Computing a work-rest schedule that minimizes fatigue

To demonstrate the benefits of our model, which is suitable for derivative-based optimization, we adopt the example of Wood et al. [74], who compute a work-rest schedule that minimizes fatigue during a grip task.

Wood et al. [74] define fatigue as the decay in maximal grip force at the end of the task. To keep their example simple, the physiological workload, the force, and the temporal pattern of the 60 sec work-rest cycles were held constant. Furthermore, the contraction intensity was bounded between 0.16 ≤ *u*_abs_ ≤ 0.48. Via grid search, they found the optimal parameters for the work-rest cycle to be a contraction lasting 22 sec at an intensity of *u*_abs_ = 0.26 followed by 38 sec of rest. The left part of Figure 7 shows the model response obtained by simulating the calibrated model for this scenario. After 60 min, MVIC force is reduced to 79.28 %.

**Figure 7:**
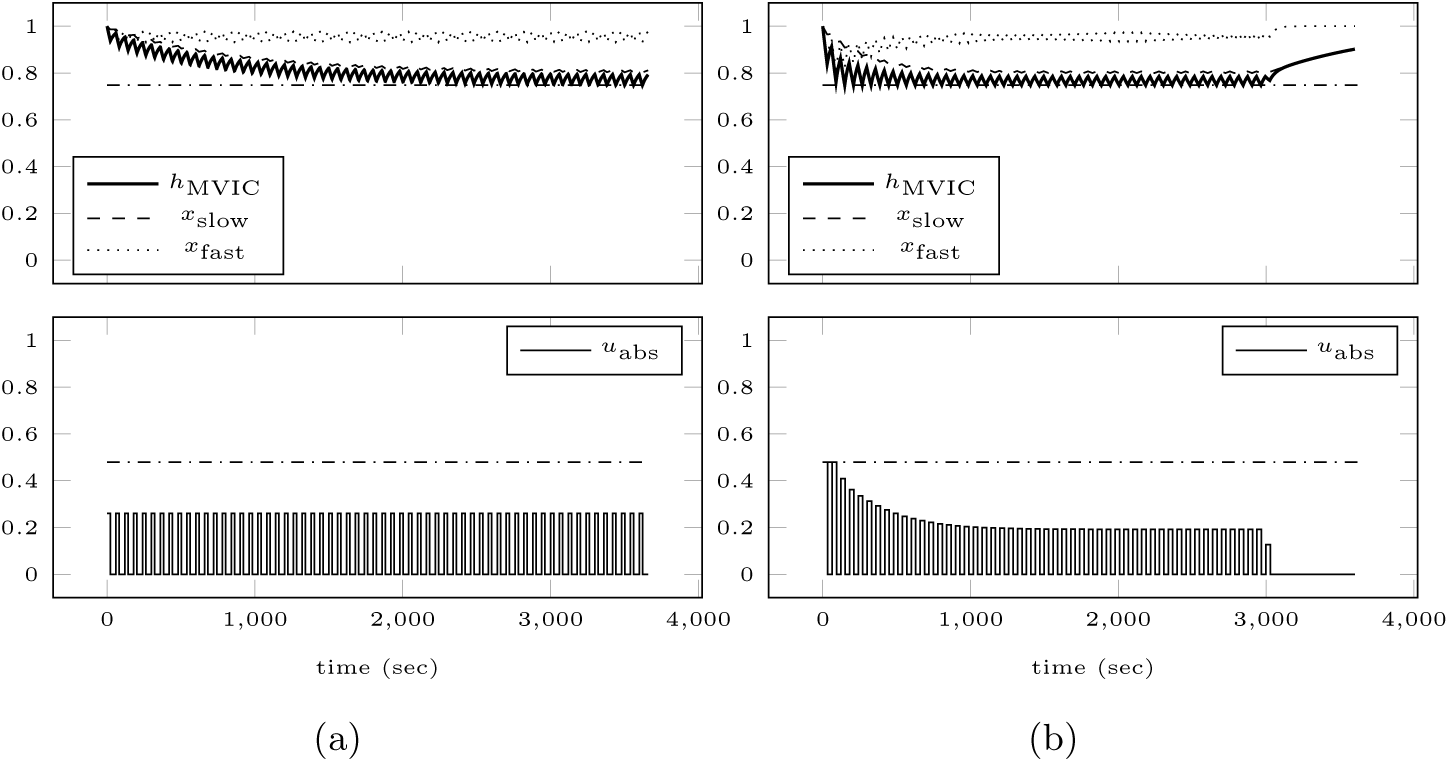
Model response obtained by simulating the calibrated model for the scenario adopted from Wood et al. [74] (a) and for an optimal loading scheme that minimizes fatigue with respect to certain safety requirements (b). The top row shows the model response and the absolute force input is illustrated in the bottom row. The horizontal dash-dotted line in the top row depicts the chosen constraint on MVIC force, whereas in the bottom row it depicts the maximum allowed external load. Using the optimal loading scheme, fatigue at the end of the task is less than when using the adopted scheme.

We now compute a work-rest schedule that minimizes fatigue for the elbow flexors. We strive to reach the same FTI of 343.20 during *T* = 60 min. Here, we allow changes of the temporal pattern and the force of the 60 work-rest cycles. Additionally, we add several safety requirements. First, MVIC force should not fall below 74.81 % of baseline, which is the minimum value obtained when simulating the scenario motivated by Wood et al. [74]. Second, *u*_abs_ should not exceed 0.48, which is the upper limit used by Wood et al. [74]. Third, to obtain a real-life feasible work schedule, the maximum duration of a contraction should not exceed 30 sec and at least 30 sec should separate two contractions.

Remembering that *x*_slow_*x*_fast_ = *h*_MVIC_ and setting *x* = (*x*_slow_, *x*_fast_, *x*_FTI_)^*T*^, the corresponding multi-stage optimal control problem [42] then reads as

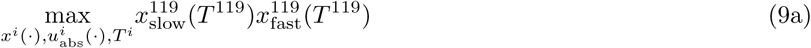

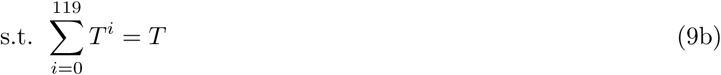

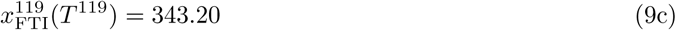

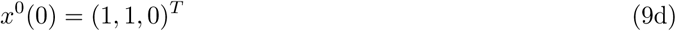

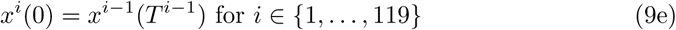

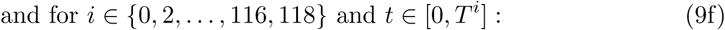

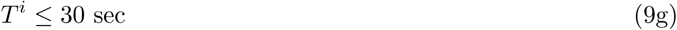

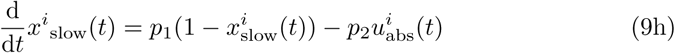

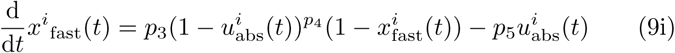

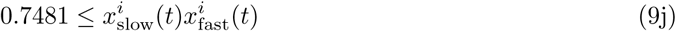

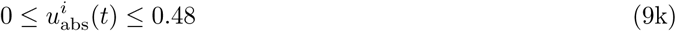

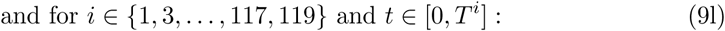

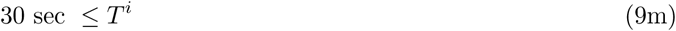

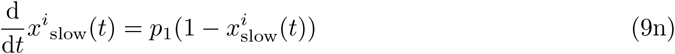

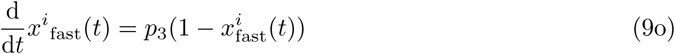

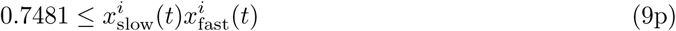

The superscript *i* denotes the model stage index. During even stages, contractions with *u*_abs_ ≤ 0.48 are possible. Odd stages are considered rest periods. The length *T*^*i*^ of each stage is optimized.

To solve the problem, we again employ a first-discretize-then-optimize strategy. The optimal control software MUSCOD-II [42, 43], which originated from the work of Bock and Plitt [8], implements a direct multiple shooting approach and is used throughout the rest of this section.

The right part of Figure 7 shows the result of the optimization. Using high intensity contractions in the beginning allows to end the task earlier and yields an MVIC force of 90.19 % of baseline after 60 min. This is an improvement of 10.91 % compared to the results obtained by adopting the scenario of Wood et al. [74] for the elbow flexors. To minimize fatigue at the end of the task, it is beneficial to accumulate work in the beginning. This can be done by either prolonging the contractions or increasing their intensity. Thus, during most of the task, at least one of the Constraints (9g), (9j), (9k), (9m), and (9p) is active. Note that, although the plots might suggest otherwise, all constraints are satisfied during the whole task.

Problem (9) contains initial conditions, endpoint constraints, control constraints, and state constraints and was chosen as a representative of a very general class of optimal control problems which can be solved with this approach. This emphasizes the advantages of an algorithmic design of loading schemes since it allows to treat even complex scenarios where intuitive planning would fail.

### 6.3 Designing an optimal isometric RT session

As a second example, we optimize an isometric RT session with the goal to increase MVIC strength of the elbow flexors. Major determinants of long-term adaptations include contraction duration, contraction intensity, inter-set rest, inter-repetition rest, and training volume [21]. As the American College of Sports Medicine [2] recommends high loads for increasing maximal strength, we choose to maximize the FTI accumulated above a threshold intensity of *u*_abs_ = 0.8, which we define as

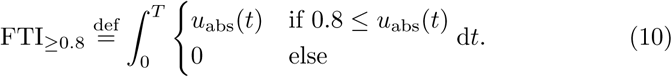

The left part of Figure 8 shows an intuitively planned isometric RT session consisting of 6 sets with 6 MVICs per set each lasting 4 sec, an inter-repetition rest of 18 sec, and an inter-set rest of 180 sec. This session is motivated by a similar one designed by Maffiuletti and Martin [49] for isometric RT of the knee extensors. However, their primary goal was not to increase strength but to examine the adaptations to different rates of force development. In contrast to the progressive increase of contraction intensity used by the authors, we choose to contract maximally during the 4 sec of a repetition, as we want to maximize the FTI_≥0.8_. Following this plan, we can accumulate an FTI_≥0.8_ of 125.0.

**Figure 8:**
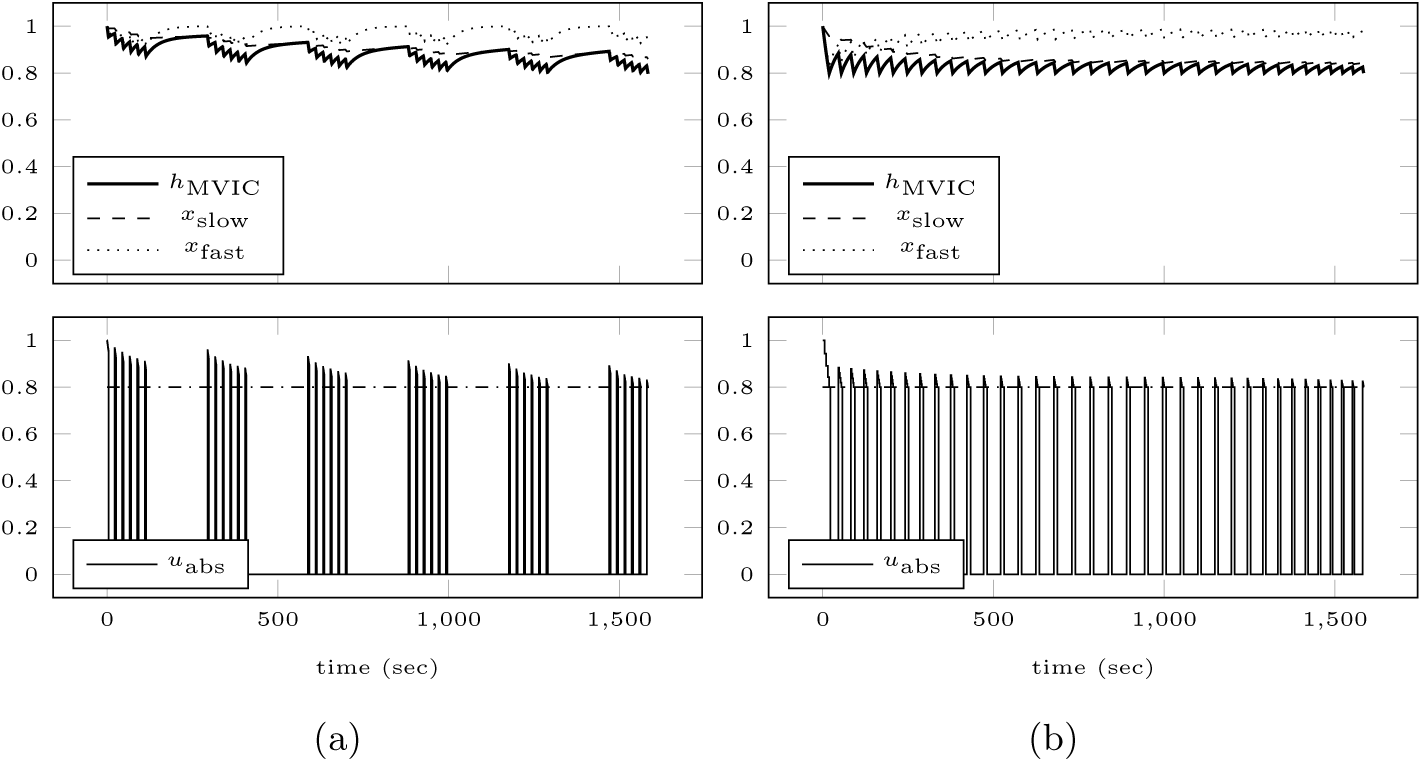
Model response obtained by simulating the calibrated model for the scenario motivated by Maffiuletti and Martin [49] (a) and for an optimal isometric RT session (b). The top row shows the model response and the absolute force input is illustrated in the bottom row. The horizontal dash-dotted line in the bottom row depicts the chosen intensity threshold. Using the optimal training plan results in a higher FTI_≥0.8_ than when using the intuitive plan.

We now formulate and solve an optimal control problem which maximizes FTI_≥0.8_. To allow a fair comparison to the adopted training plan, we limit the total time to *T* = 1584 sec and the number of repetitions to 36. Setting *x* = (*x*_slow_, *x*_fast_), the corresponding multi-stage optimal control problem [42] then reads as

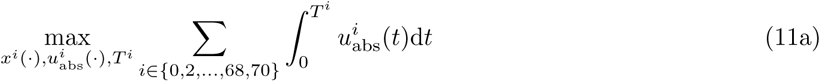

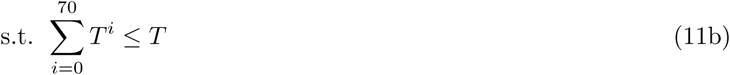

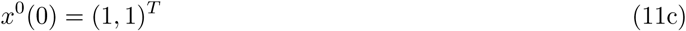

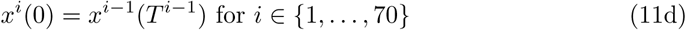

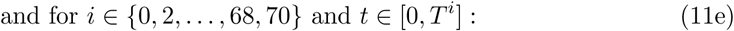

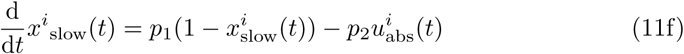

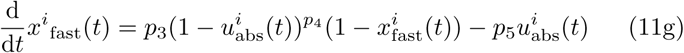

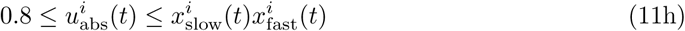

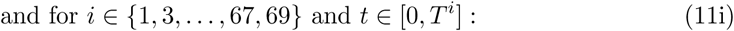

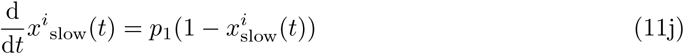

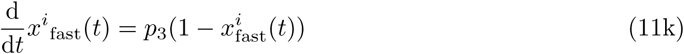

The superscript *i* denotes the model stage index. During even stages, contractions with 0.8 ≤ *u*_abs_ are possible. Odd stages are considered rest periods. The length *T*^*i*^ of each stage is optimized.

The right part of Figure 8 depicts the result of the optimization. Instead of intermittent contractions grouped into sets, the solution consists of 36 single MVICs of decreasing duration and slightly varying inter-set rests. Here, we can accumulate an FTI_≥0.8_ of 172.0. Thus, we could increase the FTI_≥0.8_ by 41 %. Since all proposed contractions are maximal, the upper part of Constraint (11h) is active during all of them. The lower part of Constraint (11h) is active at the end of each contraction. Our results would therefore support the design of isometric RT sessions without the grouping of repetitions into sets, if the goal is to maximize the FTI above a certain threshold. This was for example done by Carolan and Cafarelli [12], who employed 30 single MVICs each lasting 3 – 4 sec separated by more than 30 sec to strengthen the knee extensors.

Naturally, the improvement and outcome depends highly on the chosen intuitive setting and objective functional. Problem (11) was chosen as a further representative of complex real-life problems which can be solved with this approach. Depending on the individual goals, a broad variety of optimal control problems could be formulated.

## 7 Conclusion

Based on the model of Freund and Takala [22], we developed a simple and predictive ODE model of the time course of MVIC force which accounts for the fast and slow dynamics observed during fatiguing contractions and subsequent recovery [13]. The model was validated with a comprehensive set of published data from the elbow flexors [25, 69, 70, 66, 65] and showed promising results. Once suitable data becomes available, it will be interesting to see how the model performs for submaximal intermittent contractions, for other muscle groups, and for individual subjects.

A special focus of this work was to keep the model suitable for derivative-based optimization methods. The model can readily be employed to optimize complex loading schemes, e.g. work shifts or resistance training plans. We demonstrated this by computing a work-rest schedule that minimizes fatigue and an optimal isometric RT session as examples.

To realize the full potential of the new model, suitable optimal control problems have to be developed and solved, e.g. to minimize the risk for work-related injuries or to maximize the benefits of resistance training. For the latter, current recommendations from the sports sciences about ‘optimal’ training with respect to maximizing strength, hypertrophy, power, or local muscular endurance have to be formulated mathematically, as we demonstrated exemplary in Section 6.3. Furthermore, to facilitate fitting the model to individual persons and muscle groups, optimal experiments should be designed. For this purpose, available methods [39] can be used. Together with an extension to the more commonly used dynamic contractions, these tasks will be the topics of future research.

## Acknowledgments

We gratefully thank Dr. Janet L. Taylor of Neuroscience Research Australia, Sydney, Australia for providing and explaining the experimental data of the studies Taylor et al. [69], Taylor et al. [70], Søgaard et al. [66], and Smith et al. [65].

## Author contributions

JLH and CK conceived the idea for this work. JLH conducted the literature research, developed the model, performed the numerical experiments, and drafted the manuscript. JLH, CK, and JPS discussed and edited the draft. JLH, CK, and JPS approved the final version of the manuscript.

## Supplementary material

A Python script for simulation of the proposed model is available alongside this article on bioRxiv.org.

